# Metadichol® induced expression of circadian clock transcription factors in human fibroblasts

**DOI:** 10.1101/2024.02.21.581403

**Authors:** P.R. Raghavan

## Abstract

The circadian clock genes play a significant role in various aspects of health and disease, including sleep, metabolism, inflammation, and cancer. CRY1, BMAL1, PER1, PPARGC1A, and CLOCK are key circadian clock genes that govern the daily rhythms of diverse physiological processes in mammals. These genes create a feedback loop called a transcriptional-translational feedback loop (TTFL). In this loop, the heterodimer of CLOCK and BMAL1 stimulates the production of CRY1, PER1, and other clock genes. Moreover, the CRY1 and PER1 proteins suppressed the activity of CLOCK-BMAL1. PPARGC1A is a transcription factor that regulates the expression of clock genes and metabolic genes. Only a limited number of agonists are currently known to activate these genes. In this study, we showed that treatment with metadichol, a nanoemulsion of long-chain alcohols, significantly increased the expression of the CRY1, CLOCK, and PPARGC1A genes by 4-5-fold in human fibroblasts. Additionally, 100 ng of metadichol maintained the expression of the PER1 and BMAL1 genes, as confirmed by quantitative real-time polymerase chain reaction (Q-rt-PCR) data. The lack of significant changes in PER1 and BMAL1 expression suggested that metadichol does not directly affect these genes. However, since these genes are part of the core circadian clock machinery, their normal functioning is crucial for maintaining the circadian rhythm.

## INTRODUCTION

Circadian rhythms are intrinsic 24-hour cycles that are fundamental to physiological and behavioral processes in almost all organisms, including humans. These rhythms are driven by a biological clock, a complex orchestration of genes and proteins that regulate the timing of various bodily functions (1,2). The importance of circadian rhythms extends beyond the mere alternation of sleep and wakefulness; they are pivotal in maintaining homeostasis and overall health.

### Biological clock and circadian rhythms

The biological clock is a system of proteins encoded by clock genes that turn on and off in a precise sequence, thereby generating oscillations over a approximately 24-hour period (1). In humans, the master clock is located in the suprachiasmatic nucleus (SCN) of the brain and coordinates all the biological clocks in the body, synchronizing them to environmental cues such as light and darkness (1,2). This synchronization is crucial for the proper timing of physiological processes, including metabolism, immune response, and hormone secretion.

### Transcription factors in circadian rhythms

At the molecular level, circadian rhythms are regulated by transcription factors through positive and negative feedback loops. Key clock genes such as BMAL1 (basic helix-loop-helix ARNT like 1), CLOCK (clock circadian regulator), CRY1 (cryptochrome circadian regulator 1), and PER1 (period circadian regulator 1) play central roles in this process, as does PPARGC1A (PPARG coactivator 1 alpha). These genes control the transcription and translation of various proteins that accumulate and degrade in a rhythmic pattern, thus maintaining the circadian cycle (2). The expression of these core clock genes has far-reaching effects on cellular signaling pathways, influencing a wide array of physiological functions.

### Circadian Rhythms and Health Implications

Disruptions in circadian rhythms can lead to adverse health outcomes. Long-term sleep deprivation or chronic misalignment of circadian rhythms can increase the risk of obesity, diabetes, mood disorders, cardiovascular diseases, and even cancer(3). Understanding the role of circadian transcription factors is therefore important not only for elucidating the fundamental workings of the biological clock but also for addressing various health issues.

Research into circadian rhythms using model organisms with similar biological clock genes to those of humans has been instrumental in uncovering the mechanisms by which these rhythms influence health and disease (3). For instance, the loss of certain circadian transcription factors has been linked to conditions such as epilepsy, highlighting the importance of these factors in maintaining normal physiological functions (4).

### Circadian Rhythms and Mental Health

The relationship between circadian rhythms and mental health is particularly noteworthy. Dysregulation of circadian genes or polymorphisms in these genes may increase susceptibility to mood disorders (5). A bidirectional relationship exists between mood disorders and circadian rhythms, with mood disorders often associated with disrupted circadian clock-controlled responses (6). This finding suggested that the circadian molecular clock is not only a marker of disease states but also a contributing factor.

These are some examples of small molecules that can regulate circadian clock genes. However, the exact mechanisms and impacts of these molecules on circadian biology are not fully understood, and additional research is needed to explore their effects on other clock genes and cell types. Resveratrol, a natural polyphenol, can regulate the expression of several clock genes, such as Per1, Per2, and Bmal1, in Rat-1 fibroblasts but not in human fibroblasts (7). Resveratrol is known to have various health benefits, such as antiaging, antitumor, and anti-inflammatory effects, and to modulate the circadian clock and metabolism via Sirt1 and PGC-1α.

KL001 is a synthetic small molecule that can activate CRY proteins by stabilizing them and preventing their degradation via the ubiquitin□proteasome system (8).. KL001 can lengthen the circadian period and inhibit glucagon-induced gluconeogenesis in primary hepatocytes. KL001 also has antidiabetic effects on mouse models of type 2 diabetes.

SR9009: a synthetic small molecule that can bind to and activate the nuclear receptor REV-ERBα, also known as nuclear receptor subfamily 1 group D member 1 (NR1D1), which represses the expression of Bmal1 and other clock genes. SR9009 can alter the circadian rhythm and metabolism in mice and has been proposed as a potential treatment for obesity, diabetes, and sleep disorders (9).

Here, using Q-RT–PCR, we showed that 100 ng/ml metadichol enhances the expression of the CRY1 and CLOCK genes by 4-5-fold in fibroblasts but does not significantly alter the expression of other core circadian genes, such as PER1 and BMAL1. The upregulation of CRY1 and CLOCK could strengthen the negative feedback loop of the circadian clock, potentially leading to more robust circadian rhythms.

#### Experimental

All the experiments were designed and outsourced to the commercial service provider Skanda Biolabs Ltd., Bangalore, India. The raw data are provided in the Supplemental files.

### Cell treatment

The cells were treated for 24 hr at the indicated concentrations (Table 1) in growth media without FBS.

**Table 11.**
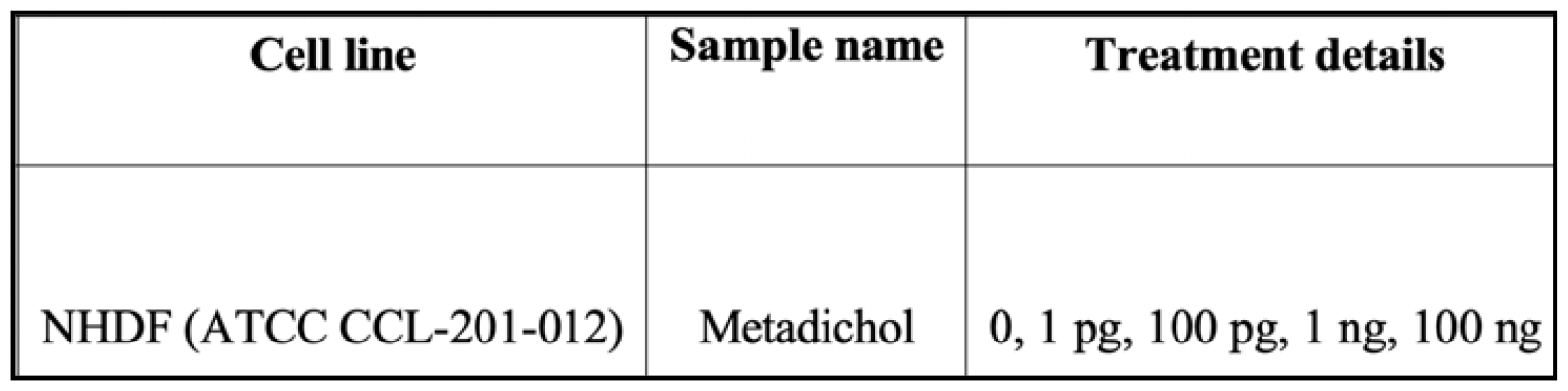

| Cell line | Sample name | Treatment details |
| --- | --- | --- |
| NHDF (ATCC CCL-201-012) | Metadichol | 0, 1 pg, 100 pg, 1 ng, 100 ng |

### Sample preparation and RNA isolation

Treated cells were harvested, rinsed with sterile 1X PBS and centrifuged. The supernatant was decanted, and 0.4 ml of TRIzol was added and gently mixed by inversion for 1 min. The samples were allowed to stand for 10 min at room temperature. To this mixture, 0.25 ml of chloroform was added per 0.4 ml of TRIzol used. He contents were vortexed for 15 seconds. The tube was allowed to stand at room temperature for 5 mins. The resulting mixture was centrifuged at 12,000 rpm for 15 min at 4°C. The upper aqueous phase was collected in a new sterile microcentrifuge tube, to which 0.5 ml of isopropanol was added, and the contents were gently mixed by inverting for 30 seconds and incubating at -20°C for 20 minutes. The contents were centrifuged at 12,000 rpm for 10 minutes at 4°C. The supernatant was discarded, and the RNA pellet was washed by adding 0.5 ml of 70% ethanol. The RNA mixture was centrifuged at 12,000 rpm at 4°C. The supernatant was carefully discarded, and the pellet was air-dried. The RNA pellet was then resuspended in 20 μl of DEPC-treated water. The total RNA yield (Table 2) was quantified using a SpectraDrop (Spectraramax i3x, Molecular Devices, USA).

**Table 2.**
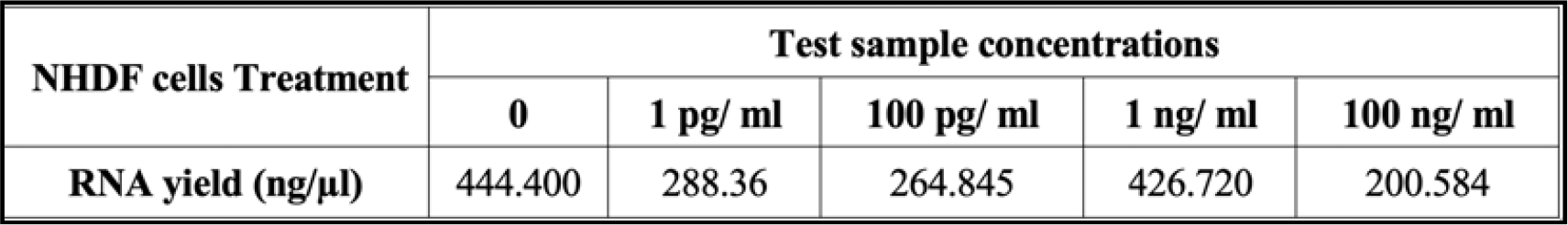

#### RT□qPCR analysis cDNA synthesis

cDNA was synthesized from 500 ng of RNA using a cDNA synthesis kit from the Prime Script RT reagent kit (TAKARA) with oligo dT primers according to the manufacturer s instructions. The reaction volume was set to 20 μl, and cDNA synthesis was performed at 50°C for 30 min, followed by RT inactivation at 85°C for 5 min using the applied biosystems Veritii. The cDNA was further subjected to real-time PCR analysis.

#### Primers and RT□qPCR analysis

The PCR mixture (final volume of 20 µl) contained 2 µl of cDNA, 10 µl of SYBR Green Master mix and 1 µl of the respective complementary forward and reverse primers specific for the respective target genes. The reaction was carried out with enzyme activation at 95°C for 2 minutes, followed by a 2-step reaction with initial denaturation and annealing at 95°C for 5 seconds and annealing for 30 seconds at the appropriate temperature for 39 cycles of amplification followed by secondary denaturation. The primer sequences are shown in Table 3.

**Table 3.**
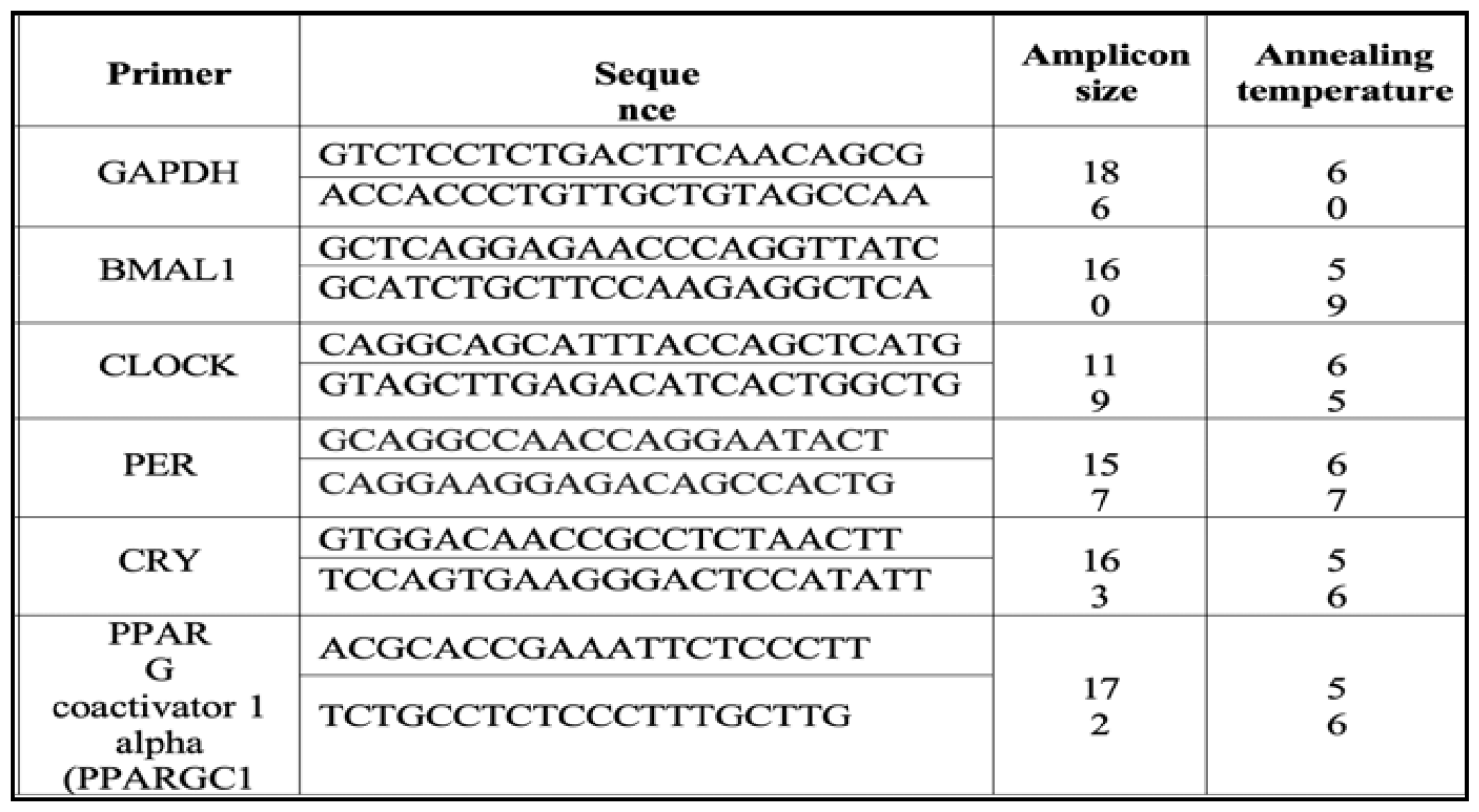

The temperature was 95°C for 5 seconds, with 1 cycle of melt curve capture ranging from 65°C to 95°C for 5 sec each. The obtained results were analyzed, and the fold change in expression or regulation was calculated.

### Results of RT□qPCR

## Discussion

The results in Figure 1 show increased expression of all the CircDIAN GENES@100 ng of Metadichol. The BMAL1 and PEr1 are maintained at normal levels where as PPARGc1A, CRY1 and CLOCK show increased expression. The circadian clock is regulated by a complex network of genes and proteins. Among the key players in this process are CLOCK, CRY1, PER1, BMAL1, and PPARGc1A..

**Figure 1.**
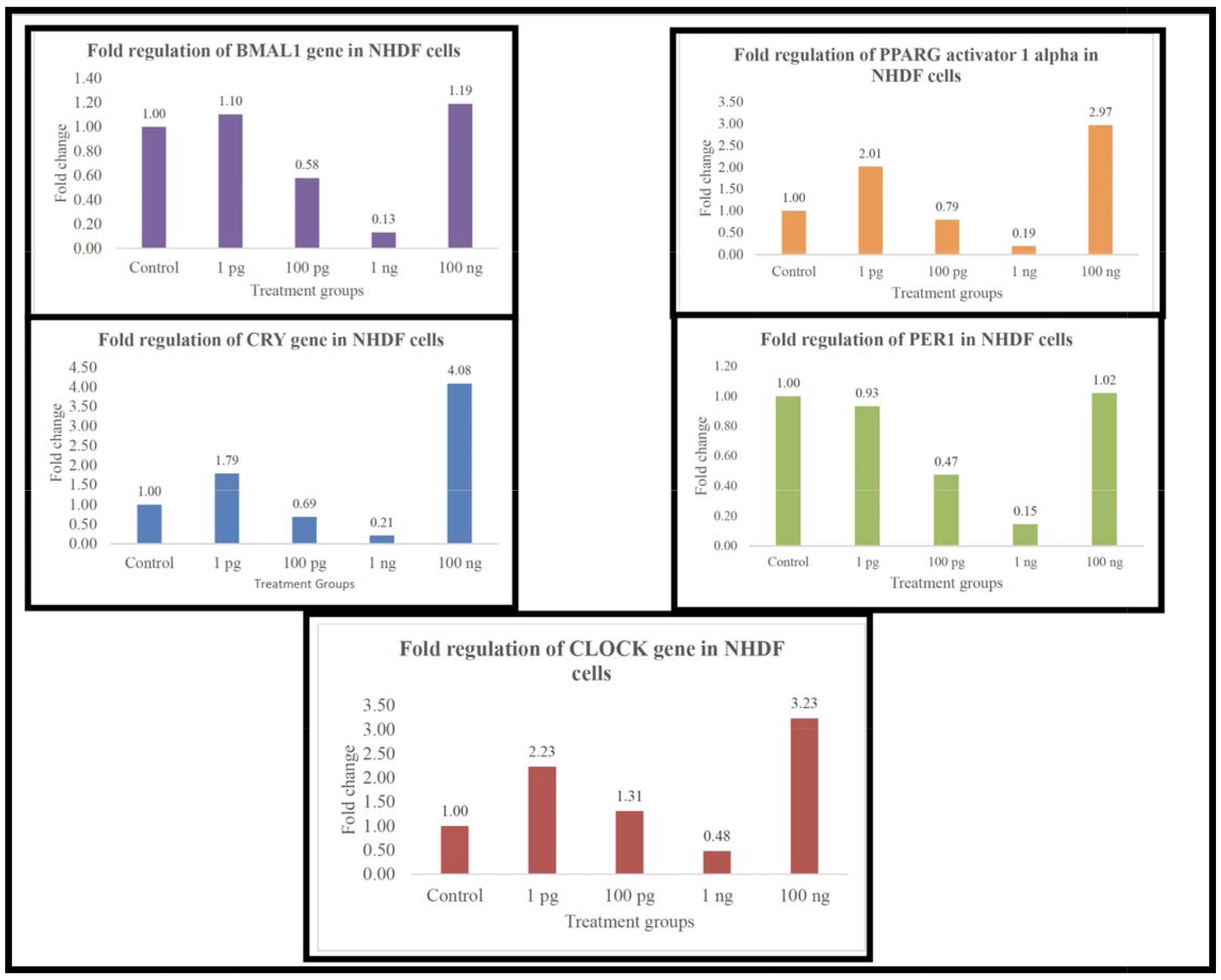
Fold regulation of circadian transcription factor expression

The circadian clock genes form a transcriptional-translational feedback loop in which the transcription f ctors CLOCK and BMAL1 activate the expression of the genes CRY1 and PER1, which encode proteins that inhibit the activity of CLOCK-BMAL1 (10,11). Other transcription factors, such as PPARGC1A and BMAL1, modulate the expression of clock genes and metabolic genes. The network formed by the transcription factors was generated using Pathway Studio (12-14) and is shown in Figure 2. The other key pathways and their impacts on diseases and organs are shown in Tables 4, 5, and 6. Circadian rhythm transcription factors are integral to the proper functioning of the biological clock and, by extension, to human health. They regulate the timing of various biological processes, and disruption of these processes can lead to a plethora of health issues.

**Table 4.** Top Pathways; Biological Processene set Clock, Per1, CRY1, Bmal1, Ppargc1a.

| Name | Parent Folder | # of Entities | Expanded # of Entities | Overlap | Percent Overlap | Overlapping Entities | p value |
| --- | --- | --- | --- | --- | --- | --- | --- |
| Circadian Clock Genes Mutations Cause Insomnia | Aging Related Diseases; Sleep Dysregulation; Sleep Dysregulation (Neurological Diseases); nerve tissue; nervous system | 26 | 41 | 4 | 9 | CRY1, CLOCK, PER1, ARNTL | 6.34966E-10 |
| Circadian Clock and Sleep Regulation by Melatonin | Amino Acid Derived Hormones; Sleep Regulation; Sleep Regulation (Biological Process); nerve tissue; nervous system | 26 | 41 | 4 | 9 | CRY1, CLOCK, PER1, ARNTL | 6.34966E-10 |
| Circadian Clock in Sleep Regulation | Sleep Regulation; Sleep Regulation (Biological Process); nerve tissue; nervous system | 32 | 55 | 4 | 7 | CLOCK, CRY1, PER1, ARNTL | 2.13638E-9 |
| Circadian Clock Genes in Suprachiasmatic Nuclei Neurons |  | 64 | 125 | 4 | 3 | CLOCK, CRY1, ARNTL, PER1 | 6.04168E-8 |
| SIRT1 Signaling in Aging | Genomic Instability Associated with Aging; Genomic Instability Associated with Aging (Aging Biology); generic; generic; nucleus | 64 | 170 | 3 | 1 | CLOCK, ARNTL, PPARGC1A | 2.90853E-5 |
| SIRT4 Signaling in Aging | Genomic Instability Associated with Aging; Genomic Instability Associated with Aging (Aging Biology); generic; generic; nucleus | 26 | 31 | 1 | 3 | PPARGC1A | 1.31176E-2 |
| Vitamin D Activates Transcription | Vitamins Biology; Vitamins Biology (Biological Process); generic; generic | 16 | 55 | 1 | 1 | CLOCK | 2.31784E-2 |
| Agomelatine Antidepressant Action | Sleep Regulation; Sleep Regulation (Biological Process); nerve tissue; nervous system | 26 | 85 | 1 | 1 | PER1 | 3.56387E-2 |
| SIRT7 Signaling in Aging | Genomic Instability Associated with Aging; Genomic Instability Associated with Aging (Aging Biology); generic; generic; nucleus | 27 | 98 | 1 | 1 | PPARGC1A | 4.09986E-2 |
| TGFB-\|COX2 | Corneal stroma remodeling | 47 | 115 | 1 | 0 | PPARGC1A | 4.79716E-2 |

**Table 5.** Organs expressed molecules enriched in the input gene set Clock,CRy1,PER1,BMAL1,PPARGc1A.

| Name | Total # of Neighbors | Gene Set Seed | Overlap | Percent Overlap | Overlapping Entities | p value |
| --- | --- | --- | --- | --- | --- | --- |
| <u>Protein entities related to testis</u> | 10374 | testis | 5 | 0 | ARNTL, PPARGC1A, CRY1, PER1, CLOCK | 0.00000E+00 |
| <u>Protein entities related to cerebral cortex</u> | 8874 | cerebral cortex | 5 | 0 | ARNTL, PPARGC1A, CRY1, PER1, CLOCK | 0.00000E+00 |
| <u>Protein entities related to brain</u> | 11768 | brain | 5 | 0 | ARNTL, PPARGC1A, CRY1, PER1, CLOCK | 0.00000E+00 |
| <u>Protein entities related to neuron</u> | 9335 | neuron | 5 | 0 | ARNTL, PPARGC1A, CRY1, PER1, CLOCK | 0.00000E+00 |
| <u>Protein entities related to lung</u> | 10040 | lung | 5 | 0 | ARNTL, PPARGC1A, CRY1, PER1, CLOCK | 0.00000E+00 |
| <u>Protein entities related to kidney</u> | 10904 | kidney | 5 | 0 | ARNTL, PPARGC1A, CRY1, PER1, CLOCK | 0.00000E+00 |
| <u>Protein entities related to liver</u> | 9996 | liver | 4 | 0 | ARNTL, CRY1, PER1, CLOCK | 0.00000E+00 |
| <u>Protein entities related to GT1-7</u> | 277 | GT1-7 | 5 | 1 | ARNTL, PPARGC1A, CRY1, PER1, CLOCK | 3.37236E-08 |
| <u>Protein entities related to hypophysis pars tuberalis</u> | 111 | hypophysis pars tuberalis | 4 | 3 | ARNTL, CRY1, PER1, CLOCK | 1.30955E-07 |
| <u>Protein entities related to suprachiasmatic nucleus neuron</u> | 154 | suprachiasmatic nucleus neuron | 4 | 2 | ARNTL, CRY1, PER1, CLOCK | 4.90711E-07 |

**Table 6.** Diseases regulated by entities enriched in the input data.

| Circadian genes<br>BMAL1 PPARGC1A,<br>CRY1, PER1, CLOCK |  |  |  |  |  |  |
| --- | --- | --- | --- | --- | --- | --- |
| Name | Total # of<br>Neighbors | Gene Set Seed | Overlap | Percent<br>Overlap | Overlapping<br>Entities | p value |
| Protein regulators of<br>glucose intolerance | 1111 | glucose<br>intolerance | 5 | 0 | BMAL1<br>PPARGC1A,<br>CRY1, PER1,<br>CLOCK | 1.97088E-07 |
| Protein regulators of<br>elevated blood pressure | 430 | elevated blood<br>pressure | 4 | 0 | BMAL1PPARG<br>C1A, CRY1,<br>CLOCK | 4.74904E-07 |
| Protein regulators of<br>hyperglycemia | 1325 | hyperglycemia | 5 | 0 | BMAL1PPARG<br>C1A, CRY1,<br>PER1, CLOCK | 4.76213E-07 |
| Protein regulators of<br>circadian rhythm sleep<br>disorder | 6 | circadian<br>rhythm sleep<br>disorder | 2 | 28 | CRY1, CLOCK | 5.06905E-07 |
| Protein regulators of<br>intermittent hypoxia | 142 | intermittent<br>hypoxia | 3 | 2 | BMAL1, CRY1,<br>PER1 | 1.93132E-06 |
| Protein regulators of<br>seasonal affective disorder | 14 | seasonal<br>affective<br>disorder | 2 | 13 | BMAL1,<br>CLOCK | 3.07320E-06 |
| Protein regulators of<br>astrogliosis | 731 | astrogliosis | 4 | 0 | BMAL1,<br>PPARGC1A,<br>CRY1, CLOCK | 3.94934E-06 |
| Protein regulators of<br>disorder of initiating and<br>maintaining sleep | 18 | disorder of<br>initiating and<br>maintaining<br>sleep | 2 | 10 | PPARGC1A,<br>CLOCK | 5.16533E-06 |
| Protein regulators of<br>sarcopenia | 213 | sarcopenia | 3 | 1 | BMAL1<br>PPARGC1A,<br>CLOCK | 6.53589E-06 |
| Protein regulators of<br>premature aging | 221 | premature<br>aging | 3 | 1 | BMAL1,<br>PPARGC1A,<br>CLOCK | 7.30046E-06 |

**Table 7.**
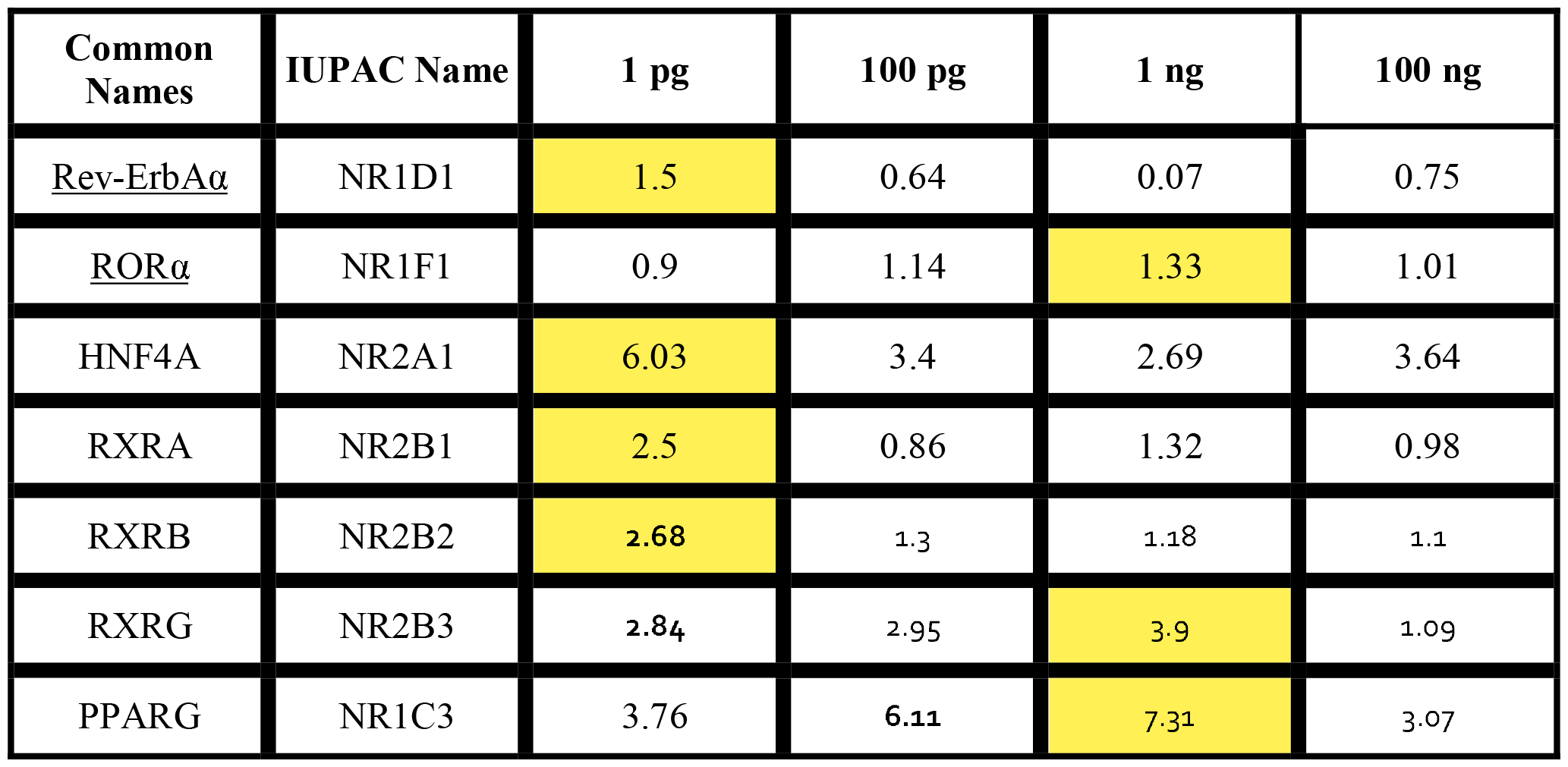
Metadichol Nuclear receptor Fold increase in Fibroblasts.

| Common Names | IUPAC Name | 1 pg | 100 pg | 1 ng | 100 ng |
| --- | --- | --- | --- | --- | --- |
| <u>Rev-ErbAα</u> | NR1D1 | 1.5 | 0.64 | 0.07 | 0.75 |
| <u>RORα</u> | NR1F1 | 0.9 | 1.14 | 1.33 | 1.01 |
| HNF4A | NR2A1 | 6.03 | 3.4 | 2.69 | 3.64 |
| RXRA | NR2B1 | 2.5 | 0.86 | 1.32 | 0.98 |
| RXRB | NR2B2 | 2.68 | 1.3 | 1.18 | 1.1 |
| RXRG | NR2B3 | 2.84 | 2.95 | 3.9 | 1.09 |
| PPARG | NR1C3 | 3.76 | 6.11 | 7.31 | 3.07 |

**Table 8.** Metadichol and Sirt1 and Sirt6-fold expression.

| Gene | 1 pg | 100 pg | 1 ng | 100 ng | Control |
| --- | --- | --- | --- | --- | --- |
| Sirt1 | 3.29 | 2.30 | 3.48 | 6.92 | 1.00 |
| Sirt6 | 2.19 | 1.97 | 3.01 | 4.26 | 1.00 |

**Figure 2.**
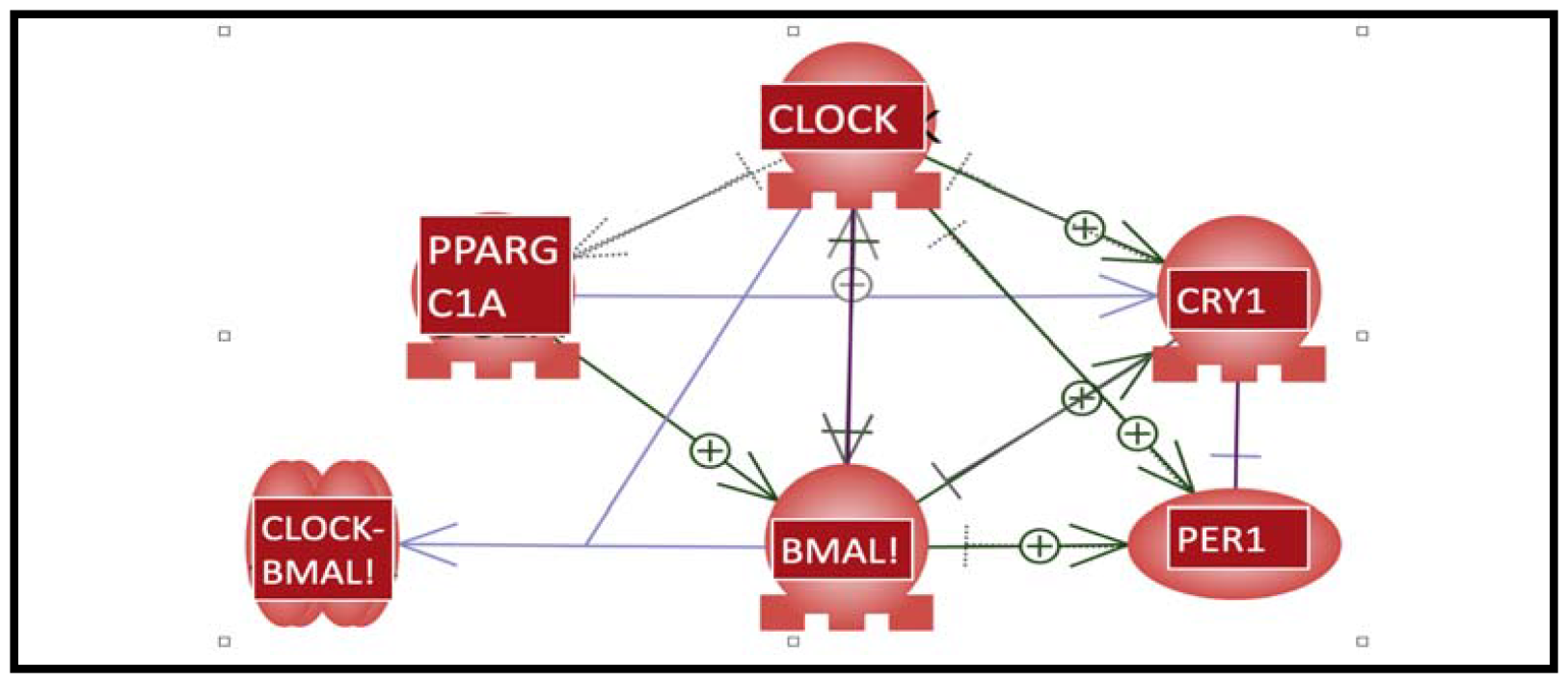
Network of the Circadian gene network.

### Interactions

The interactions and roles of CLOCK, CRY1, PER1, BMAL1, and PPARGc1A in regulating the circadian rhythm are as follows:

### CLOCK and BMAL1

CLOCK and BMAL1 form a heterodimer that activates the transcription of other clock genes, including the PER1, CRY1 and cryptochrome (Cry) genes. This activation occurs by binding to E-box promoter elements in the genome, thus activating a large number of clock-controlled genes (15). This activation leads to the production of the PER and CRY proteins, which accumulate and form complexes that repress their own transcription by binding to the CLOCK:BMAL1 complex, creating a negative feedback loop. This cycle repeats approximately every 24 hours, contributing to the circadian rhythm (15-17).

### PER1 and CRY1

PER1 and CRY1 form a complex that inhibits the activity of CLOCK-BMAL1, creating a negative feedback loop. This inhibition leads to the suppression of their own transcription, thus forming the negative arm of the circadian clock (18). Along with other PER and CRY proteins, PER1 is involved in the regulation of the expression of genes outside of the core clock genes. It has been suggested that PER1 and CRY1/2 can regulate the expression of genes outside of the core clock, indicating a broader role in cellular function (19).

The expression levels of PER1 and BMAL1 are maintained through a complex interplay of transcriptional, posttranscriptional, and posttranslational modifications. Disruptions in the circadian rhythms of PER1 and BMAL1 have been linked to various physiological and pathological conditions. For instance, circadian misalignment and disruptions in circadian rhythms have been associated with an increased risk of various diseases, including cancer (20-23). The maintenance of normal levels of PER1 and BMAL1 is crucial for proper functioning of the circadian rhythm, and disruptions in their expression can have significant implications for human health (24). Mrtadichol@100 ng was used to maintain normal levels.functioning of the circadian rhythm, and disruptions in their expression can have significant implications for human health (). Mrtadichol@100 ng was used to maintain normal levels.

### PPARGc1A

PPARgC1A is a transcriptional coactivator that plays a critical role in the maintenance of energy metabolism. It has been found to interact with a gene regulatory network of the circadian clock (25). PGC-1alpha stimulates the expression of clock genes, notably Bmal1 (Arntl) and Rev-erbalpha (Nr1d1), through coactivation of the ROR family of orphan nuclear receptors (26). PPARgC1A, BMAL1, and CLOCK are interconnected in the regulation of circadian rhythm and metabolic processes. BMAL1 and CLOCK have been identified as significant upstream regulators, and they function as heterodimers (BMAL1:CLOCK) that bind the enhancer box (E-box) regulatory element, regulating the transcription of Period (PER) and Cryptochrome (CRY) genes, which form the negative arm of the circadian clock (27).

### Metadichol and Circadian gene expression

It is highly unlikely that Metadichol can directly activate all these transcription factors. It is through upstream genes that this happens. Previously, we showed that metadichol induces the expression of all 48 nuclear receptors. (28) and the sirtuin (1-7) genes (29). Sirt1 and Sirt6 play important roles in maintaining circadian rhythm (30-31). RXr, RORa, NR1F1, NR1D1 (REV-ERBa) and HNF4A. The fold changes in nuclear receptor expression in Fibrobalst cells induced by metadichol are shown in Table 2.

The expression of the CLOCK gene is regulated (32) by the nuclear receptor RORα (retinoic acid receptor-related orphan receptor alpha) and the transcriptional coregulator receptor interacting protein 140 (RIP140). RORα is involved in a secondary feedback loop that regulates the CLOCK gene, and RIP140 was identified as a modulator of CLOCK that participates in a feedback mechanism affecting the circadian clock.

Additionally, the nuclear receptor nuclear receptor subfamily 1 group D member 1 (NR1D1) is involved in regulating the expression of target genes, including the CLOCK gene. Therefore, both RORα and NR1D1 are responsible for the expression of the CLOCK gene (33.).

The nuclear receptor responsible for the expression of the Cry1 gene is HNF4A (34). HNF4A (NR2A1) is critical for circadian rhythm maintenance and period regulation in liver and colon cells. It acts differently from other CLOCK:BMAL1 repressors and is able to bind CRY proteins, thereby inhibiting CLOCK:BMAL1 through a mechanism independent of CRY1. On the other hand, the nuclear receptor REV-ERBα, also known as NR1D1, is involved in the regulation (35) of the BMAL1 gene, which is part of the core circadian clock. While REV-ERBα regulates the expression of BMAL1, it does not directly regulate the expression of the cry1 gene (36). Therefore, HNF4A is the nuclear receptor responsible for the expression of the cry1 gene.

The nuclear receptor responsible for the expression of the BMAL1 gene is the retinoid-related orphan receptor α (RORA), also known as NR1F1. RORA stimulates transcription from the BMAL1 promoter, thereby regulating the expression of the BMAL1 gene (37). This regulation is part of the complex network of interactions involved in the circadian clock, where RORA plays a key role in activating the transcription of clock genes, including BMAL1. Therefore, RORA is the nuclear receptor responsible for the expression of the BMAL1 gene.

The nuclear receptors responsible for the expression of the PPARGC1A (PGC-1α) **<u>g</u>**ene are the retinoid X receptor (RXR) and peroxisome proliferator-activated receptor gamma (PPAR-γ). PGC-1α binds to PPAR-γ and coactivates PPAR-γ to stimulate the transcription of genes involved in various metabolic processes, including mitochondrial biogenesis and energy metabolism (38). The interaction between PGC-1α and PPAR-γ is essential for the regulation of genes related to energy metabolism and other physiological functions. Therefore, PPAR-γ and its coactivator PGC-1α play crucial roles in the regulation of various metabolic pathways. Thus, **the action of metadichol** on nuclear **receptors** (39) leads to **decreased** expression of **circadian** genes.

### Sirtuins 1 and 6 and their role in and circadian processes

Metadichol also **induces Sirtuin expression in fibroblasts** (28). The fold increases **are** shown in Table ()

Sirtuins are a family of NAD+-dependent protein deacetylases that play crucial roles in various cellular processes, including aging, metabolism, and the regulation of circadian rhythms. Sirtuins, particularly SIRT1, have been shown to directly interact with components of the circadian clock machinery, influencing the regulation of circadian rhythms (40).

- SIRT1 and the CLOCK-BMAL1 complex: SIRT1 has been found to bind to the CLOCK-BMAL1 complex, a core component of the circadian clock, and promote the deacetylation of BMAL1. This interaction is crucial for the proper functioning of the circadian machinery. SIRT1 activity is regulated in a circadian manner, suggesting a feedback mechanism in which SIRT1 modulates the circadian clock, which in turn influences SIRT1 activity..
- Regulation of PER2: SIRT1 also regulates the stability of PER2, another core clock protein, through deacetylation This modification affects the stability and degradation of PER2, further illustrating the role of SIRT1 in modulating circadian rhythms (41).
- Aging and Circadian Control: The function of SIRT1 in the brain, particularly in the suprachiasmatic nucleus (SCN) — the central circadian clock in mammals — has been shown to decrease with aging. This decrease in SIRT1 activity may contribute to the aging-related deterioration of circadian function(42).
- Metabolic Link: SIRT1 mediates the connection between the circadian clock and cellular metabolism through its dependency on NAD+, a cofactor whose levels oscillate in a circadian manner. This relationship underscores the role of SIRT1 in aligning metabolic processes with the circadian cycle (43).

#### Other Sirtuins

While SIRT1 is the most studied sirtuin in the context of circadian rhythms, other members of the sirtuin family also contribute to circadian regulation. SIRT6 has been identified as a critical regulator of circadian transcription, suggesting that it also serves as an interface between the circadian clock and cellular metabolism (44).

#### Implications for Health and Disease

The involvement of sirtuins in circadian regulation has significant implications for understanding the pathophysiology of various diseases, including metabolic disorders, aging-related diseases, and cancer. Disruptions in circadian rhythms have been linked to a range of health issues, and the role of sirtuins in this process highlights their potential as therapeutic targets (45-58).

In summary, sirtuins, particularly SIRT1 and SIRT6, play a critical role in the regulation of circadian rhythms through their interactions with core clock components and modulation of metabolic processes. Their activity influences the stability and function of circadian proteins, aligning physiological processes with the environmental day-night cycle. This intricate relationship between sirtuins and the circadian clock underscores the importance of these proteins in maintaining cellular and organismal homeostasis.

## Conclusions

Metadichol activation of these transcription factors is associated with neuronal function (59) and development, may have implications for the treatment of circadian rhythm-related disorders, such as sleep phase disorders, and could also play a role in the regulation of pluripotent stem cell differentiation, offering avenues for regenerative medicine.

Ongoing research is crucial for further understanding the complex interactions between these transcription factors and the myriad of physiological pathways they influence. This knowledge holds the potential for developing novel therapeutic strategies for circadian-related disorders and improving overall human health. Additionally, the finding that metadichol, a small molecule, induces the expression of transcription factors such as Clock, CRY1, PER1, BMAL1, PPARGC1A, and ARNTL is relevant for several reasons. 100 ng of Metadichol enhanced CRY1 and CLOCK gene expression by 4-5-fold in fibroblasts, while the changes in PER1 and BMAL1 expression were similar to those in the control group; these findings could have significant implications for the circadian biology network. The lack of significant changes in PER1 and BMAL1 expression suggested that metadichol does not directly affect these genes. However, since these genes are part of the core circadian clock machinery, their normal functioning is crucial for maintaining the circadian rhythm. Gene expression induced by Metadichol could enhance the robustness of the circadian rhythm in fibroblasts.

Additionally, Metadichol has been found to modulate the expression of genes involved in viral infections (60) and inhibit the entry of SARS-CoV-2 into host cells, indicating its potential role in antiviral responses. Moreover, the ability of metadichol to induce the expression of critical transcription factors such as Yamanaka factors (61) in various cell types holds promise for its application in stem cell therapy and reprogramming of somatic cells. The advantage of this approach is that metadichol is nontoxic (62-64), well tested and commercially available for a decade. These findings collectively suggest that the ability of Metadichol to influence the expression of various transcription factors has significant potential for diverse applications in neurology, virology, and regenerative medicine.

## Supporting information

raw Q-RT-PCR data

## Data availability

All the raw data are presented in the manuscript and supplementary materials.

## Supplementary material

Raw Data; Q-RT□PCR

