## Supplementary material for "Metadichol® induced expression of circadian clock transcription factors in human fibroblasts": raw Q-RT-PCR data

### Study Report

#### *In vitro* evaluation of circadian rhythm family genes in NHDF cells

### *In vitro* evaluation of circadian rhythm family genes in NHDF cells

**Cellular Marker**

1) BMAL1
2) CLOCK
3) PER
4) CRY
5) PPARG coactivator 1 alpha

**Chemicals/ reagents used**

1] Sterile workplace, tips, pipettes, centrifuge tubes, 6-well plate etc.
2] RPMI or DMEM Medium
3] Fetal bovine serum (FBS)
4] 0.05% trypsin
5] RNase free environment or workstation
6] Autoclaved deionised water
7] TRIzol reagent (Life technologies, Cat. No. 15596026)
8] Chloroform
9] Chilled isopropanol
10] Chilled 70% ethanol
11] Ice cold Phosphate buffered saline (PBS, pH 7.4)
12] Autoclaved, oven dried 1.5 and 2 ml micro-centrifuge tubes and PCR tubes
13] cDNA synthesis kit or Prime script RT Reagent kit (TAKARA, Cat. No. RR037A)
14] SyBR Green I (PCR Biosystems, Cat. No. PB20.15)
15] Dimethyl Sulphoxide (DMSO)
16] Target specific forward and reverse primer etc.

**Cell line and culture condition details**

| Sr. No. | Cell line | Media details |
|---|---|---|
| 1 | NHDF (ATCC CCL-201-012) | DMEM media + 10% FBS + 1% Penicillin-streptomycin |

**Cell treatment**

Cells were treated for 24 hrs at the following concentrations using growth media without FBS.

**Table 1: Treatment concentration**

| Sr. No. | Cell line | Sample name | Treatment details |
|---|---|---|---|
| 1 | NHDF (ATCC CCL-201-012) | Metadichol | 0, 1 pg, 100 pg, 1 ng, 100 ng |

**Sample Preparation and RNA Isolation**

Treated cells were harvested and rinsed with sterile 1X PBS and centrifuged. The supernatant was decanted and 0.4 ml of TRIzol was added and gently mixed by inversion for 1 min. Samples were allowed to stand for 10 minutes at room temperature. To this 0.25 ml chloroform was added per 0.4 ml of TRIzol used. The contents were vortexed for 15 seconds. The tube was allowed to stand at room temperature for 5 mins. The resulting mixture was centrifuged at 12,000 rpm for 15 mins at 4˚C. Upper aqueous phase was collected to a new sterile micro centrifuge tube to which 0.5 ml of isopropanol was added and gently mixed by inverting the contents for 30 seconds and incubated at -20˚C for 20 minutes. The contents were centrifuged at 12,000 rpm for 10 minutes at 4˚C. Supernatant was discarded and the RNA pellet was washed by adding 0.5 ml of 70% ethanol. The RNA mixture was centrifuged at 12,000 rpm at 4˚C. Supernatant was carefully discarded and the pellet was air dried. The RNA pellet was then re-suspended in 20 µl of DEPC treated water. Total RNA yield was quantified using Spectra drop (Spectramax i3x, Molecular devices, USA)

**Table 2: Total RNA yield**

| NHDF cells Treatment | Test sample concentrations | | | | |
|---|---|---|---|---|---|
| | **0** | **1 pg/ ml** | **100 pg/ ml** | **1 ng/ ml** | **100 ng/ ml** |
| **RNA yield (ng/µl)** | 444.400 | 288.36 | 264.845 | 426.720 | 200.584 |

#### qPCR analysis

##### cDNA synthesis

The cDNA was synthesized from 500 ng of RNA using the cDNA synthesis kit from Prime script RT reagent kit (TAKARA) with oligo dT primer according to the manufacturer's instructions. The reaction volume was set to 20 µl and cDNA synthesis was performed at 50°C for 30 min, followed by RT inactivation at 85°C for 5 min using applied bio-systems, Veritii. The cDNA was further used for real time PCR analysis.

##### Primers and qPCR analysis

The PCR mixture (final volume of 20 µl) contained 2 µl of cDNA, 10 µL of SyBr green Master mix and 1 µM of respective complementary forward and reverse primers specific for respective target genes. The reaction was carried out with enzyme activation at 95°C for 2 minutes followed by 2 step reaction with initial denaturation and annealing cum extension step at 95°C for 5 seconds, annealing for 30 seconds at appropriate respective temperature amplified for 39 cycles followed by secondary denaturation at 95°C for 5 seconds, 1 cycle with melt curve capture step ranging from 65°C to 95°C for 5 secs each. The obtained results were analysed and fold expression or regulation was calculated.

**Table 3: Primer details**

| Sr. no. | Primer | Sequence | Amplicon size | Annealing temperature |
|---|---|---|---|---|
| 1 | GAPDH | GTCTCCTCTGACTTCAACAGCG<br>ACCACCCTGTTGCTGTAGCCAA | 186 | 60 |
| 2 | BMAL1 | GCTCAGGAGAACCCAGGTTATC<br>GCATCTGCTTCCAAGAGGCTCA | 160 | 59 |
| 3 | CLOCK | CAGGCAGCATTTACCAGCTCATG<br>GTAGCTTGAGACATCACTGGCTG | 119 | 65 |
| 4 | PER | GCAGGCCAACCAGGAATACT<br>CAGGAAGGAGACAGCCACTG | 157 | 67 |
| 5 | CRY | GTGGACAACCGCCTCTAACTT<br>TCCAGTGAAGGGACTCCATATT | 163 | 56 |
| 6 | PPARG coactivator 1 alpha (PPARGC1A) | ACGCACCGAAATTCTCCCTT<br>TCTGCCTCTCCCTTTGCTTG | 172 | 56 |

#### 1] GAPDH

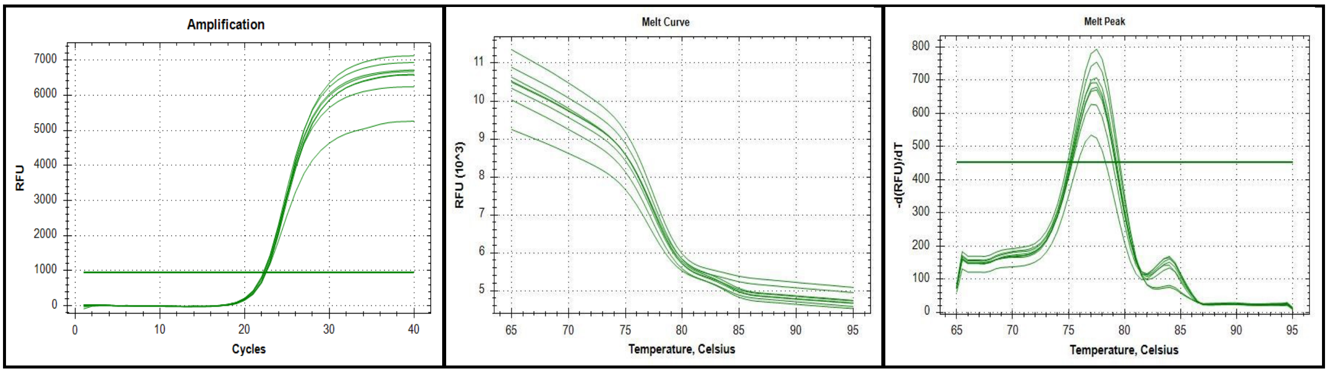


**Fig 1: Amplification curve, Melt curve and Melt peak of GAPDH gene**

#### 2] BMAL-1

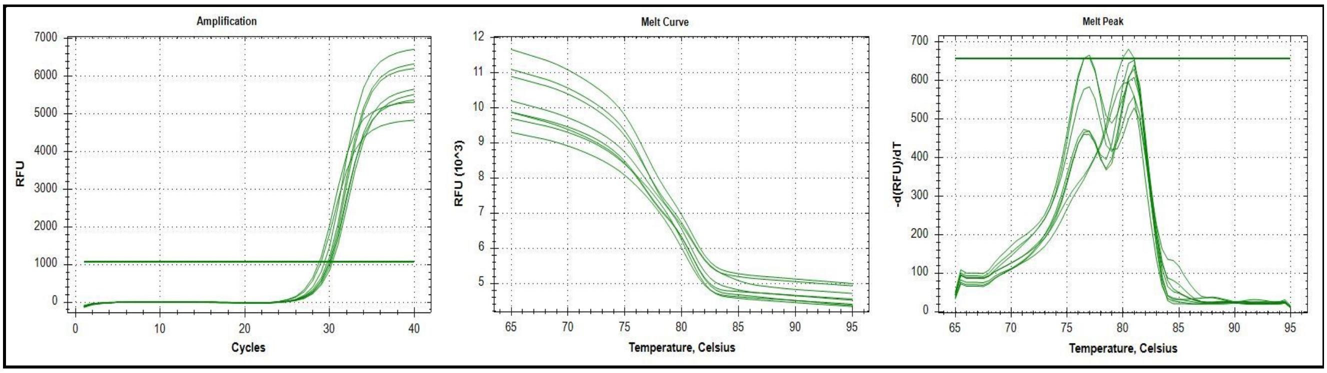


**Fig 2: Amplification curve, Melt curve and Melt peak of BMAL-1 gene**

#### 3] CLOCK

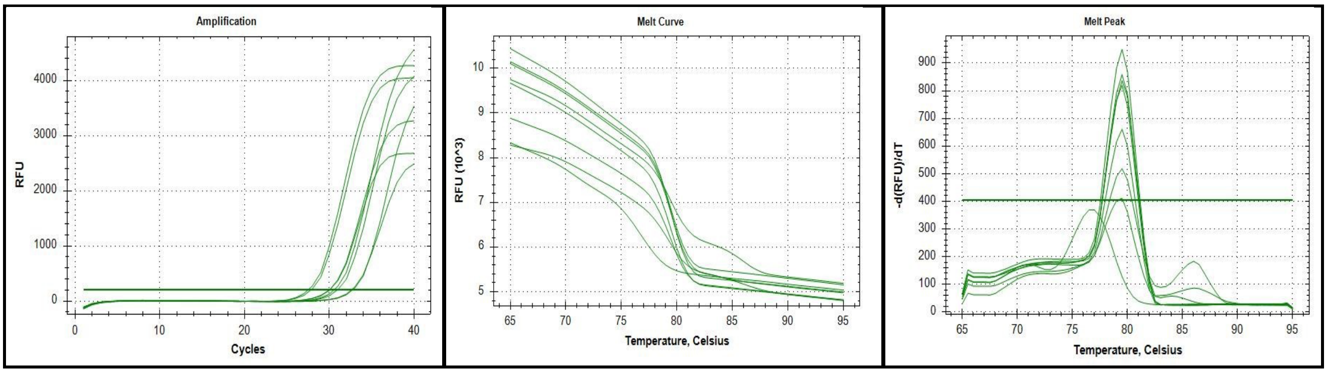


**Fig 3: Amplification curve, Melt curve and Melt peak of CLOCK gene**

4] PER

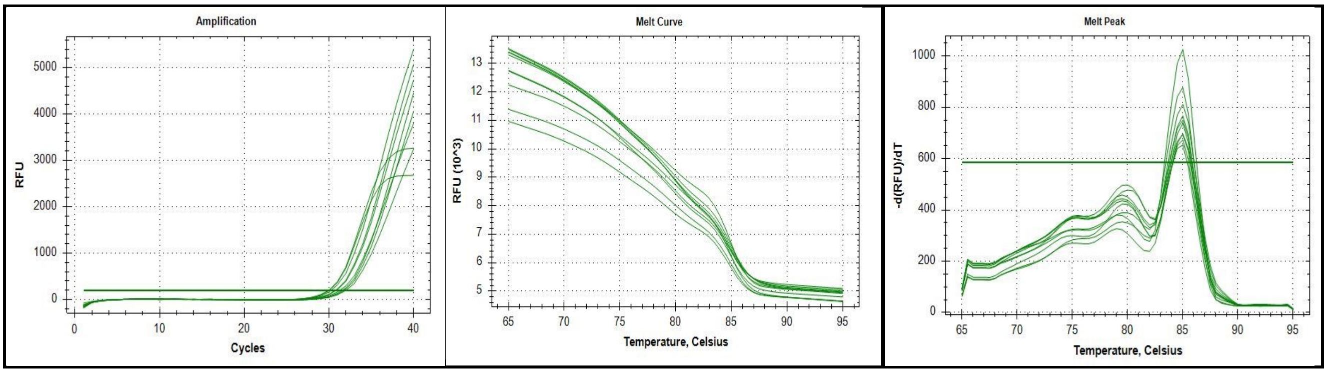


**Fig. 4: Amplification curve, Melt curve and Melt peak of PER gene**

**5] CRY**

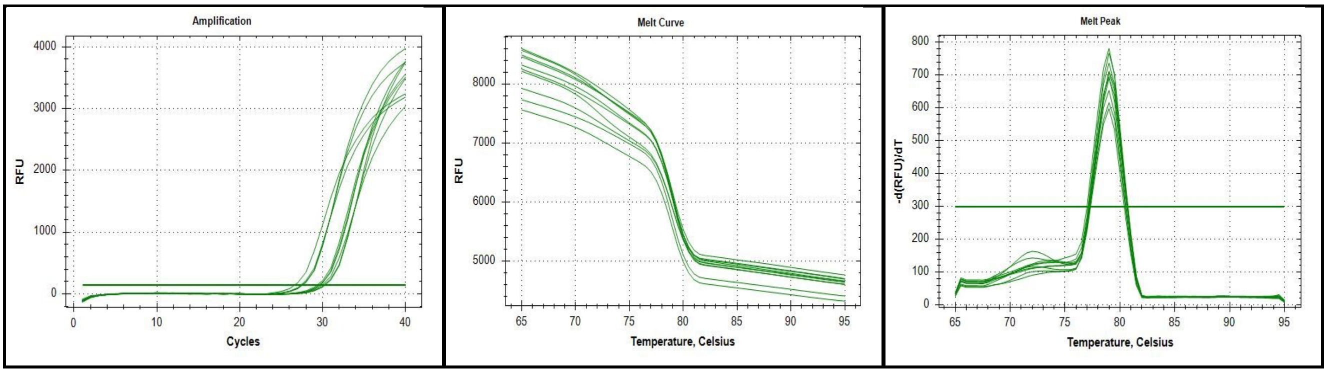


**Fig. 5: Amplification curve, Melt curve and Melt peak of CRY gene**

**6] PPARG coactivator 1 alpha**

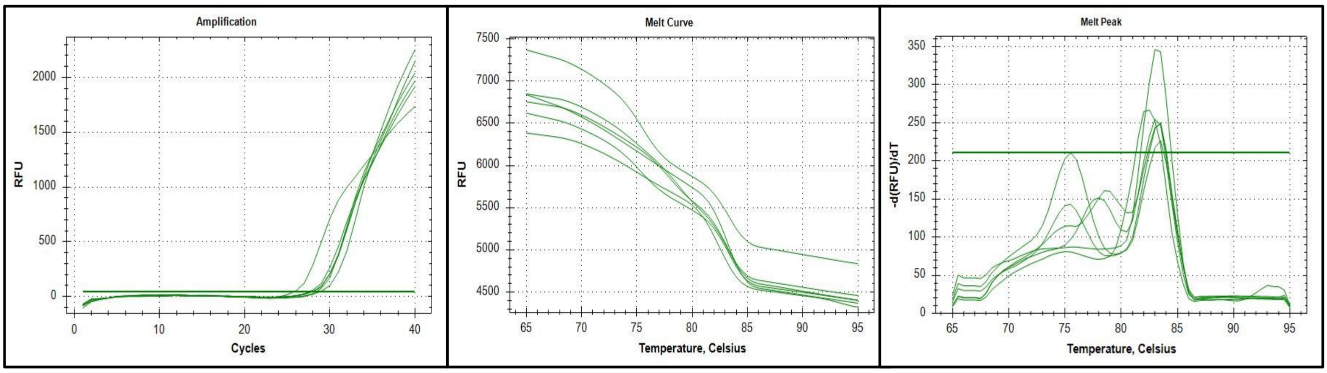


**Fig. 6: Amplification curve, Melt curve and Melt peak of PPARG coactivator 1 alpha gene**

**Table 4: Relative normalized expression of BMAL1 gene**

| Treatment Group | BMAL1 | GAPDH | ΔCq | ΔΔCq | Fold change 2^-ΔΔCq |
|---|---|---|---|---|---|
| Control | 29.61 | 20.56 | 9.05 | 0.00 | 1.00 |
| 1 pg | 29.87 | 20.96 | 8.91 | -0.14 | 1.10 |
| 100 pg | 29.61 | 19.77 | 9.84 | 0.79 | 0.58 |
| 1 ng | 29.56 | 17.58 | 11.98 | 2.93 | 0.13 |
| 100 ng | 28.33 | 19.53 | 8.80 | -0.25 | 1.19 |

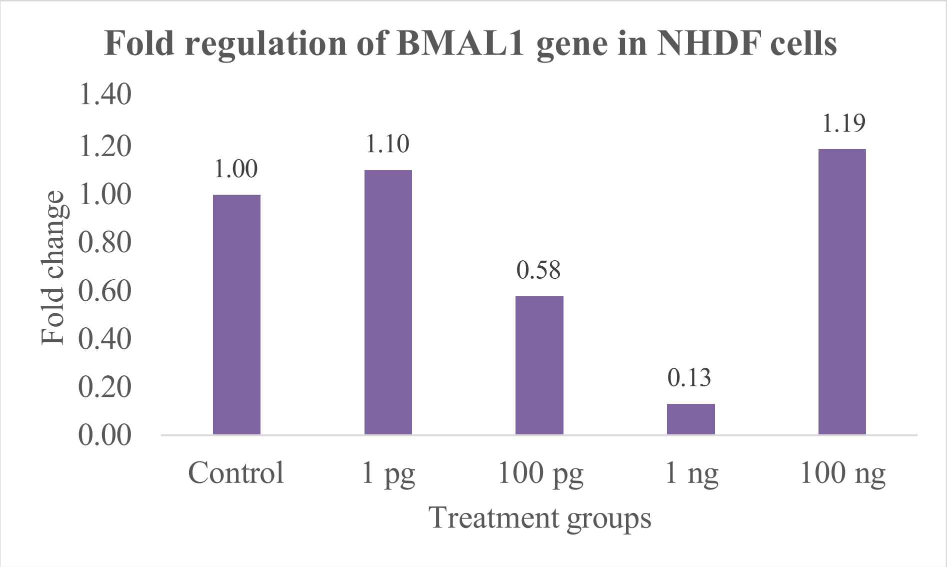


**Fig 6: Relative normalized expression of BMAL1 gene**

**Table 5: Relative normalized expression of CLOCK gene**

| Treatment Group | CLOCK | GAPDH | ΔCq | ΔΔCq | Fold change 2^-ΔΔCq |
|---|---|---|---|---|---|
| Control | 32.36 | 20.56 | 11.8 | 0.00 | 1.00 |
| 1 pg | 30.41 | 19.77 | 10.64 | -1.16 | 2.23 |
| 100 pg | 32.37 | 20.96 | 11.41 | -0.39 | 1.31 |
| 1 ng | 30.45 | 17.58 | 12.87 | 1.07 | 0.48 |
| 100 ng | 29.64 | 19.53 | 10.11 | -1.69 | 3.23 |

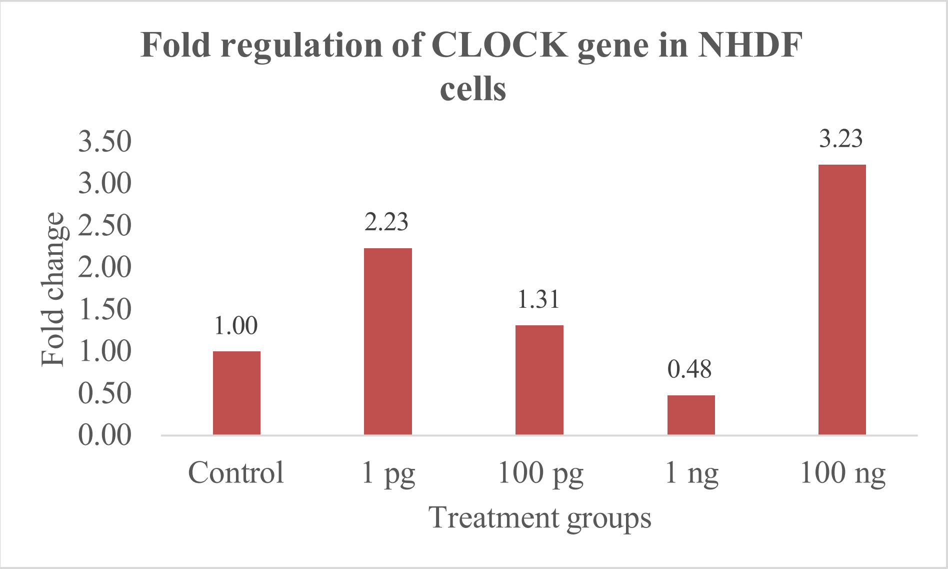


**Fig 7: Relative normalized expression of CLOCK gene**

**Table 6: Relative normalized expression of PER gene**

| Treatment Group | PER | GAPDH | ΔCq | ΔΔCq | Fold change 2^-ΔΔCq |
|---|---|---|---|---|---|
| Control | 30.32 | 20.56 | 9.76 | 0.00 | 1.00 |
| 1 pg | 30.82 | 20.96 | 9.86 | 0.10 | 0.93 |
| 100 pg | 30.61 | 19.77 | 10.84 | 1.08 | 0.47 |
| 1 ng | 30.12 | 17.58 | 12.54 | 2.78 | 0.15 |
| 100 ng | 29.26 | 19.53 | 9.73 | -0.03 | 1.02 |

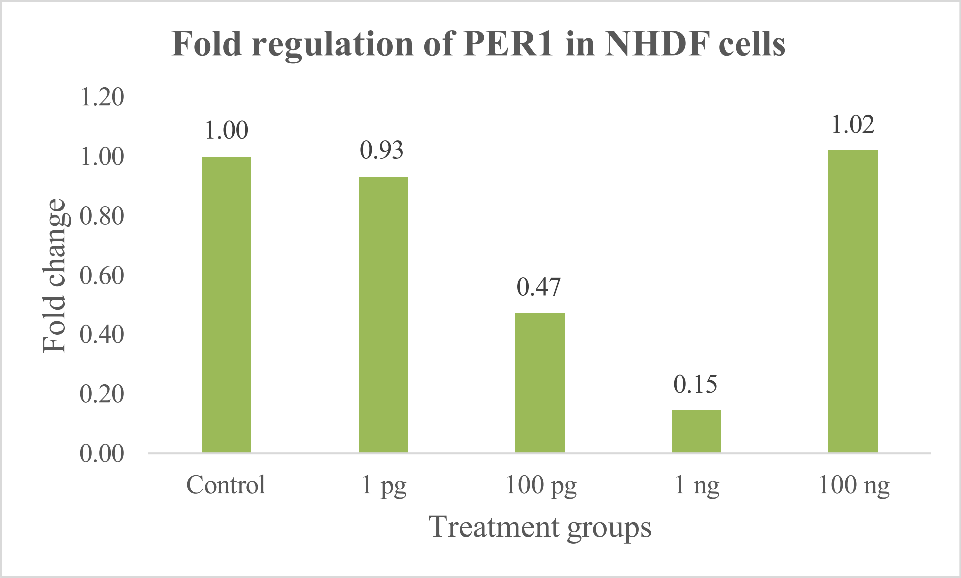


**Fig 8: Relative normalized expression of and PER gene**

**Table 7: Relative Normalised expression of CRY gene**

| Treatment Group | CRY | GAPDH | ΔCq | ΔΔCq | Fold change 2^-ΔΔCq |
|---|---|---|---|---|---|
| Control | 30.21 | 20.56 | 9.65 | 0.00 | 1.00 |
| 1 pg | 29.77 | 20.96 | 8.81 | -0.84 | 1.79 |
| 100 pg | 29.96 | 19.77 | 10.19 | 0.54 | 0.69 |
| 1 ng | 29.45 | 17.58 | 11.87 | 2.22 | 0.21 |
| 100 ng | 27.15 | 19.53 | 7.62 | -2.03 | 4.08 |

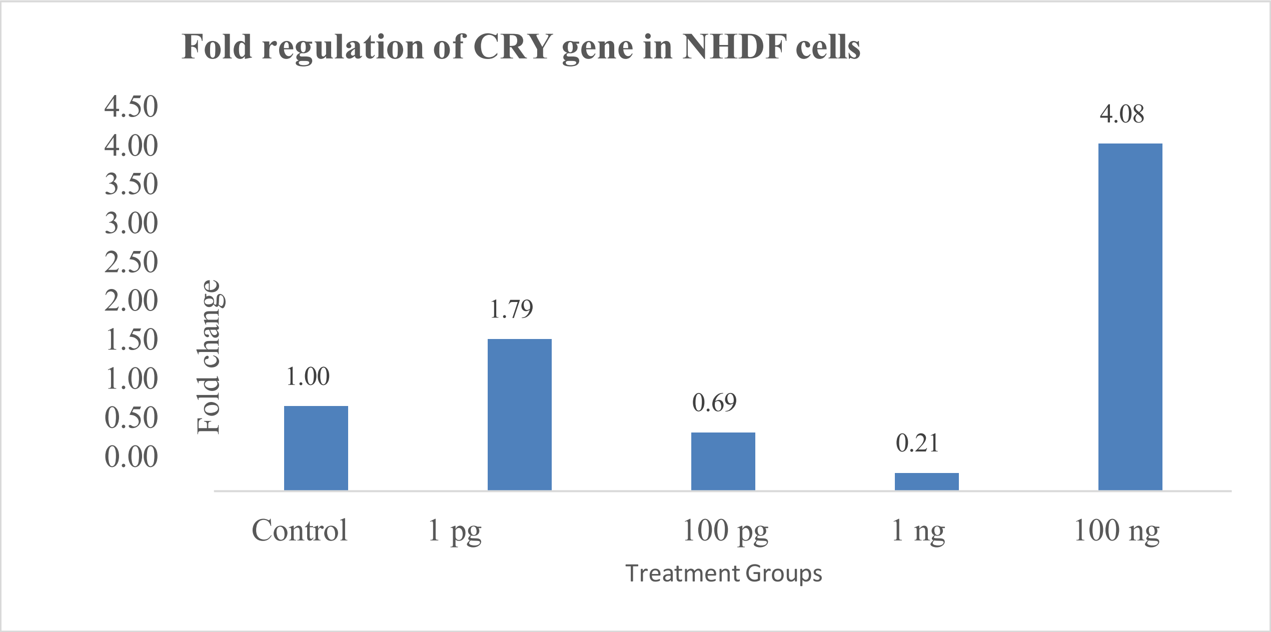


**Fig 9: Relative normalized fold expression of CRY gen**

**Table 7: Relative Normalised expression of PPARG co-activator1 Alpha gene**

| Treatment Group | PPARGC 1 Alpha | GAPDH | ΔCq | ΔΔCq | Fold change 2^-ΔΔCq |
|---|---|---|---|---|---|
| Control | 28.09 | 20.56 | 7.53 | 0.00 | 1.00 |
| 1 pg | 27.48 | 20.96 | 6.52 | -1.01 | 2.01 |
| 100 pg | 27.64 | 19.77 | 7.87 | 0.34 | 0.79 |
| 1 ng | 27.54 | 17.58 | 9.96 | 2.43 | 0.19 |
| 100 ng | 25.49 | 19.53 | 5.96 | -1.57 | 2.97 |

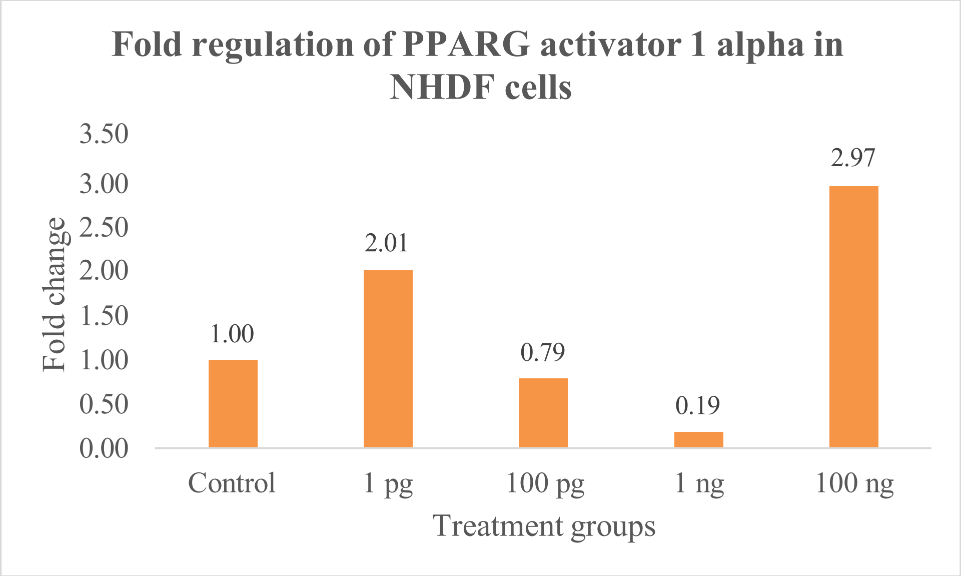


**Fig 10: Relative normalized fold expression of PPARG co-activator1 Alpha gene**

**Comments**

The RT PCR analysis of BMAL1, CLOCK, PER, CRY and PPARG coactivator 1 alpha were carried out in NHDF Cell line and fold regulation was determined. The gene expression levels of BMAL1, CLOCK, PER, CRY and PPARG coactivator 1 alpha were significantly up regulated by 1.19, 3.23, 1.02, 4.08, and 2.97 fold respectively in case of 100 ng treated cells compared to untreated cells in NHDF cells.
